# Life history stage-dependent nuclear selection in chimeric fungi

**DOI:** 10.64898/2026.04.23.718558

**Authors:** Jiayu Li, Ariel Fitzmorris, Justina Martelli, Alex Mela, N. Louise Glass, Amy S. Gladfelter, Marcus Roper

## Abstract

A single hyphal compartment of the filamentous fungus *Neurospora crassa* may contain tens or hundreds of nuclei, sharing macromolecules with each other, and, via a continuous cytoplasm, with the nuclei in other compartments. Nuclear lineages acquire mutations with each mitosis, which, combined with the autonomous mitosis of nuclei, has fueled speculation that multilevel selection may occur, both upon the mycelium, and upon individual nuclear populations. Here, we combine experiments on fungal chimera formed from two auxotrophically and epitopically labeled nuclear populations, with specially created microscopy toolkit for extracting the proportions of the two nucleotypes to analyze the strength of nuclear-level selective forces at different stages in the fungal life history. We find strong nucleotype-selective forces during spore-germination and establishment of the mycelium, and no evidence of selection on nuclei inside a growing mycelium. The kinetics of mycelial initiation from individual spores therefore allow for the selection of nuclear compositions best adapted to the fungus’ environment, in accordance with the hypothesized function of unicellular life history stages for purging deleterious mutations.

## 1 Introduction

A fungal mycelium is a single, cytoplasmically connected network that can contain potentially millions of haploid, often genetically diverse nuclei [1, 2]. Despite being bathed in a common cytoplasm, nuclei divide autonomously of each other [3, 4]. Yet, unlike multinucleate animals, in which somatic cells are segregated from the germ line [5], new nuclear lineages introduced into mature *Neurospora crassa* mycelia both disperse through the entire mycelium and are incorporated into its asexual spores [2, 6]. Most genetic diversity occurring within a mycelium arises from mutation, and sharing macromolecules across a common cytoplasm allows nearby nuclei to complement loss-of-function mutations [7], potentially allowing faster mutation rates [3, 8]. Distinctively, hyphae can also fuse and exchange nuclei. Although limited by heteokaryon compatibility loci [9], nuclear transfers between mycelia of divergent species occur frequently enough to allow horizontal gene transfer [10].

Guido Pontecorvo, a pioneer of microbial genetics, argued in a 1946 paper: “we may be justified in considering a hypha as a mass of cytoplasm with a population of nuclei.” going on to note that these nuclear populations are subject to the same processes that shape populations of organisms including “(1) variation in numbers; (2) drift - i.e., random variation in the proportions of the different kinds of nucleus; (3) migration - i.e., influx and outflow of nuclei, following hyphal anastomoses; (4) mutation; and (5) selection.” [11].

Although, in the 80 years since Pontecorvo’s lecture, the complexity of syncytism remains at the fulcrum of fungal evolution [12, 13] and behavior [14], the extent to which population genetic forces operates within the mycelium remains little understood. James et al. [15] showed that in the dikaryotic basidiomycete fungus *Heterobasidion parviporum*, proportions of two nucleotypes present within the fungus shifted in response to substrate, and that dikaryotic mycelia could grow slower or faster than homokarya containing only one nuclear species, which the authors interpreted as evidence of conflict or cooperation between the two nuclear populations present. In contrast to basidiomycete fungi, ascomycetes do not, generally, fix the number of nuclei present within each of their hyphal compartments, creating even greater possibility for nucleotypic variation [16]. Meunier et al. [17] studied the pseudohomothallic ascomycete, *Neurospora tetrasperma*, which maintains two populations of nuclei that have different mating types. In some but not all lineages, heterokarya of the two mating types showed convergence in the fraction of the two nucleotypes, which was independent of starting proportions, and also varied across life-history stages. Intriguingly, proportions of the two nucleotypes affected mycelial fitness, measured by spore production rates. In a parallel study, the authors further observed that many genes differentially expressed by the two nucleotypes showed additive levels of expression in the heterokaryon [18]; in other words, that the proportions of the two nucleotypes control the abundances of many of the mRNAs that they express. Although these data establish that changing nucleotype proportions can change expression profiles in heterokaryotic mycelia, the authors stop short of previous authors [12, 3] who argued that changing nucleotype proportions in response to a heterogeneous environment or host creates phenotypic plasticity for the mycelium.

Laboratory evolution experiments in *N. crassa* reveal the evolutionary dynamics of how new nuclear lineages emerge [19]. Propagating mycelia over many generations led to the emergence of new lineages, that when combined in heterokarya with wild type nuclei, preferentially occupied conidia. The authors called these nucleotypes *selfish* because, as they became more abundant, the mycelial fitness, measured via its spore production [20], decreased.

Pontecorvo [11] proposed an experimental test of the hypotheses underlying his theory of nuclear population genetics; heterokarya containing two or more auxotrophic nuclear populations will be complemeneted for one or both components of the population by supplying the missing nutrients externally, so “the medium, […] determines different optimal nuclear ratios”. Although we are not aware of work using auxotrophs to systematically perturb nucleotype interactions in fungal heterokarya (see, however, [3]), both engineered and natural auxotrophs are widely used to manipulate the population genetics of mononucleate microbes [21, 22], including for biosynthesis [23] and to create evolutionary games [24].

Here, we combine interactions between auxotrophically-complementary nucleotypes within *N. crassa* heterokarya with an enhanced version of a previously created image analysis pipeline for determining nuclear composition of a mycelium from fluorescent labeling of the two nuclear populations. The method offers advantages over genetic tools for measuring nucleotype proportions that require harvesting several square centimeters of mycelium, and that do not allow for finer scale analysis of nuclear proportions. In the original pipeline [25], we analyzed heterokarya built from nucleotypes without auxotrophic labels. The presence of two versions of the labeled-histone gives each nucleus within the mycelium a detectable phenotype. Due to sharing of proteins, in a heterokaryotic hypha, all nuclei will, over time, take up both fluorescent labels. However, turnover of proteins means that the fluorescence of spores eventually reflects only the nuclei they contain [6]. Our approach differs from [26], which used flow cytometry to phenotype large numbers of spores by fluorescence. Multi-channel fluorescence measurements give a presence-absence signal; i.e., indicating whether a spore contains either or both of the two nucleotypes. However, without data on the number of nuclei each spore contains, the data are insufficient to infer the proportions of the two nucleotypes.

In this study, we replicate some of the experiments from [26], using a microscopy based method similar to [25] that has been enhanced to provide statistical estimates for the genetic composition of each spore (i.e., how many nuclei of each type are present), based on its measured fluorescence. In [25], only heterokaryotic spores with two nuclei were analyzed, since the genotypes of such spores are unambiguous (i.e., a dikaryotic spore that contains both GFP and DsRed must contain one nucleus of each nucleotype). In the pipeline developed for this work, we measure the total red and green fluorescence of each heterokaryotic spore, and count the nuclei it contains. Taking these data together enables us to develop *a posteriori* estimates of how many nuclei of each nucleotype each spore contains.

Our study uses the labeled strains previously created in [26]. These strains have complementary auxotrophic deficits (Fig. 1), so that only heterokarya containing both nucleotypes can grow on unsupplemented media [26]. We seek to identify at which life history stages selective media influences nuclear ratios. In one set of experiments, we added conidial suspensions directly to plates containing either minimal media or supplemented so that one of the two nucleotypes could grow as a homokaryon. In a second set of experiments, we added the conidial suspensions onto minimal media first, forcing them to form a heterokaryotic mycelium. After 24 hours (or *≈* 1.5 cm) growth, we then transferred an agar block cut from the growing periphery of the mycelium to selective media. Our experiment exposed the heterokaryon to media selection at different life history stages: on the spores in the first experiment and on the heterokaryotic mycelium in the second. Within a heterokaryotic mycelium, proteins expressed by the two nucleotypes are shared between them, so we expect selection forces to be weak or even absent. It is common for the strength of selective forces to be life-history dependent in multicellular organisms: In many developing organisms there is an early segregation of germ and somatic cell lines, so that only mutations occurring very early in embryonic development can be transmitted intergenerationally [5].

**Figure 1:**
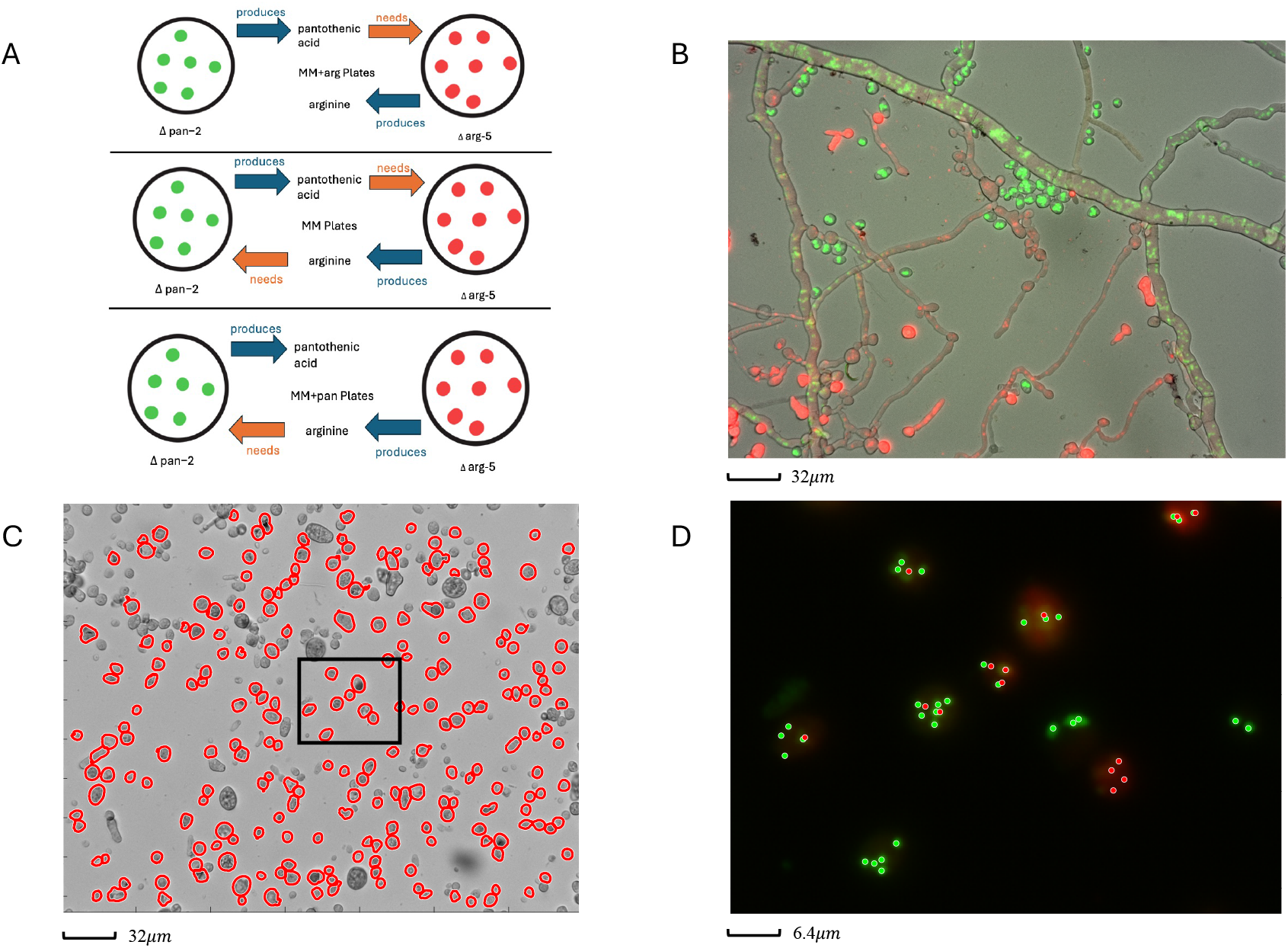
Heterokarya created from complementary auxotrophic mutants allow selective forces on nuclear populations to be measured across the fungal life history. **A:** Schematic showing (inter)dependence between *hH1-DsRed* Δ*arg-5* and *hH1-eGFP* Δ*pan-2* nuclei on selective and non-selective media. **B:** Germinating *hH1-DsRed* Δ*arg-5* spores introduced at the edge of a 20 hour post-germination *hH1-eGFP* Δ*pan-2* mycelium introduce new nuclei into the mycelium. The last two panels represent two views of the spores used to measure nucleotype proportions. **C:** transmitted light image with outlines of detected spores shown in red. Black rectangle shows an image region that is magnified in panel **D**, including nuclei above GFP thresholds (green spots) or DsRed thresholds (red spots).

## 2 Material and Methods

### 2.1 Experimental methods

In all experiments, we used previously constructed *his-3::hH1-eGFP* Δ*pan-2* [26] and *his-3::hH1-DsRed* Δ*arg-5* [27] strains of *N. crassa*. Both strains express a fluorescently tagged histone H1-protein. The Δ*pan-2* strain is unable to synthesize pantothenate (vitamin B5), whereas Δ*arg-5* is unable to synthesize the amino acid arginine. Neither strain can grow on unsupplemented media as a homokaryon, but a heterokaryon containing both nucleotypes can grow because Δ*arg-5* nuclei supply the enzymes needed to make pantothenate and Δ*pan-2* nuclei supply the enzymes needed to make arginine. All mycelia were started from asexual spores (conidia) harvested from 7-to 14-day-old cultures that were maintained on slants, containing Vogel’s minimal medium [28] supplemented with arginine (for Δ*arg-5* ) or calcium panthothenate (for Δ*pan-2* ). Spore suspensions from both strains were prepared in DI water, filtered through cheesecloth to remove hyphal fragments, and adjusted to a concentration of *≃* 8 *×*10^9^ spores/ml, and spore inocula with equal or skewed proportions of the two nucleotypes were made by mixing two homokaryotic suspensions. All mycelia were started from 5 *µ*l of spore suspension on 60 mm minimal media (Vogels MM 1 *×* plus 1.5% sucrose, 1.5% agar) plates.

Two experimental designs were used. 1. Mycelia were started by inoculating spore suspensions directly onto the selective plates. 2. Heterokaryotic mycelia were preformed by inoculating spores onto unsupplemented MM plates. 20 hours after inoculation, heterokaryotic mycelium was collected from the *≃* 3 cm diameter mycelium, by cutting approximately 3 mm *×* 3 mm squares of agar from the periphery containing its growing hyphal tips, and transferring the agar square, mycelial side down, to the center of a new plate. All experiments were replicated between 3 and 8 times.

To measure nuclear proportions, spores were harvested from around the periphery of each plate between 14-21 days post-inoculation using an applicator stick to scrape all aerial hyphae down to the agar surface, and suspended in 1-2 ml DI water, before filtering through cheesecloth to remove hyphal fragments. Spore suspensions were prepared on glass slides, sealed with petroleum jelly, and imaged under 260*×* magnification using a Zeiss AxioZoom microscope (Zeiss, White Plains, NY), under transmitted light, DsRed and GFP filtersets [25]. Around 20 microscope images were taken for each spore sample, containing *≃*5000 total spores per plate.

### 2.2 Calculation of nucleotype proportions

Nuclear composition of each spore was inferred using custom written image analysis code in MATLAB R2023b (Mathworks, Natick, MA). *a posteriori* likelihood distributions were calculated for the proportion, *p*, of Δ*arg-5* nuclei present in the imaged spores, by measuring the outline of each spore, as well as the number of nuclei, *N*_*i*_, it contains (Fig. 1C & D), followed by its total intensity in both the GFP-channel and DsRed-channel (resp., *G*_*i*_ and *R*_*i*_). By compiling data across all spores in a given plate, we construct the distribution of intensities for a single nucleus of either type. Given *N*_*i*_, we can then calculate the likelihood of our model recreating any pair of observed intensities *G*_*i*_ and *R*_*i*_. Finally we maximize the joint likelihoods of (*G*_*i*_, *R*_*i*_) over all measured spores in two parameters, *α* and *β* that characterize the likelihood of any partitioning of *N*_*i*_ into Δ*pan* and Δ*arg* nuclei. We report the likelihood maximizing values of the parameters *α* and *β*, from which we can deduce the proportions of the two nucleotypes in the heterokaryotic mycelium (see SI appendix for calculation details).

## 3 Results and Discussion

We first validated our assumption that the distribution of nuclei within each spore could be modeled via a beta-binomial distribution. First we showed that the nuclei populating each spore are not independently drawn from the underlying mycelium by analyzing spores containing either one or two nuclei. These spores can be genotyped (i.e., their fluorescence is a direct readout of the nucleotypes present). For each of 34 analyzed plates we considered ≳ 300 total spores. For each plate, under the binomial assumption, we have three fitting parameters – the total number of mono- and dikaryotic spores and the proportion, *p* of *hH1-DsRed* Δ*arg-5* nuclei, and 5 data points (i.e., the number of mono and dikaryotic spores and the proportions of *hH1-DsRed* Δ*arg-5* or *hH1-eGFP* Δ*pan-2* homokaryons or heterokaryons). Under the binomial assumption, the ratio of *hH1-DsRed* Δ*arg-5* to *hH1-eGFP* Δ*pan-2* homokaryons spores should be *p* : (1*−p*) for monokaryons and *p*^2^ : (1*−p*)^2^ for dikaryons. Fitting *p* to the data, we tested for goodness of fit to the observed distributions via a *χ*^2^ test, obtaining 𝒫 values against those ranged from 0.001 to 0.65, against independence, when *p* was estimated from all spores, and from 0.001 to 0.18 when only the monokaryotic spores were used to estimate *p*. Under either fitting method, all but 3 of the replicates had 𝒫 *<* 0.05 against the presumption of independence. See SI Fig. 1A & B.

We then compared the goodness of fit between the binomial distribution, which assumes independent nuclei, and the beta-binomial distribution, which allows for correlations between nucleotypes to be modeled, again using mono- and dikaryotic spores. Since these are not nested hypotheses, and have different numbers of fitting parameters, we compared them via their AIC scores, finding that in all but 2 of the 34 experiments, the Beta binomial distribution had a lower AIC score than the binomial distribution (SI Fig. 1C), and the mean AIC difference across all experiments was 49.7, meaning that the likelihood of seeing the observed data was approximately *e*^25^ greater for the beta binomial model than the binomial model.

Having validated our data analysis tools, we used the image analysis pipeline with best beta binomial fits to estimate the nucleotype composition of spores. First, we tested for variations of nucleotype composition across spores taken from different locations in a single mycelium. We divided the periphery of a single MM plate, that had been inoculated with a 1:1 suspension of *hH1-DsRed* Δ*arg-5* and *hH1-eGFP* Δ*pan-2* spores, into six equally sized sectors, and then calculated likelihoods (likelihood of the observed spores, given *p*) for each sector. In each sector, *p*-likelihoods were sharply peaked at the MLE, reflecting narrow uncertainty in *p*-values. The MLE values for *p* were consistent, though not identical, between sectors (Fig. 2A), consistent with previous work showing that the topology of *Neurospora*’s mycelial network and the flows of cytoplasm within the network keep nucleotypes present within the mycelium well-mixed along its growing periphery [6]. Accordingly, in the remaining experiments analyzed in this work, we sampled spores along the entire periphery of the mycelium.

**Figure 2:**
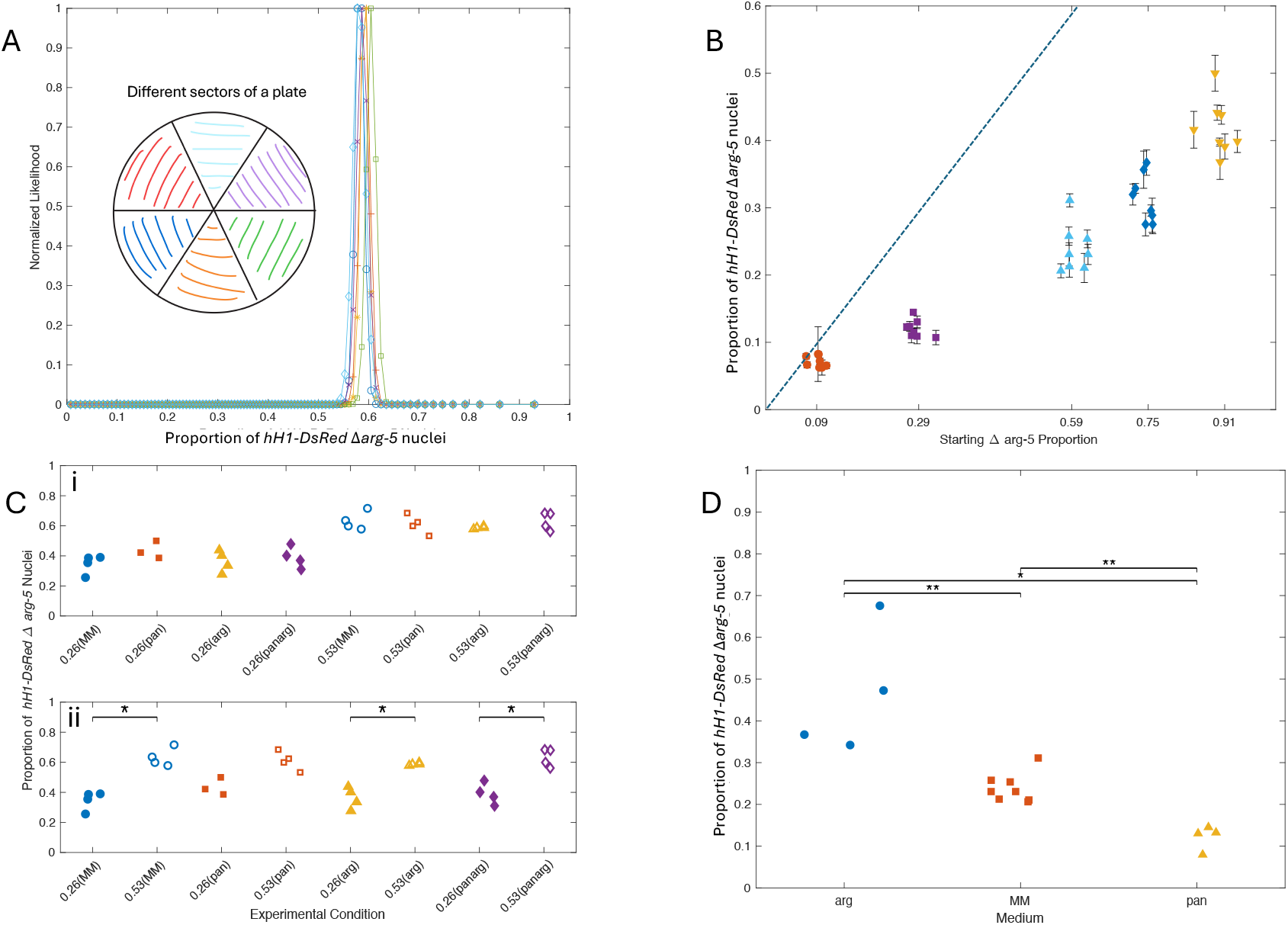
Proportion of *hH1-DsRed* Δ*arg-5* nuclei depends on initial conditions of spores, but not when formed heterokarya are grown on selective media. **A**: Normalized log-likelihood given *p* (proportion of *hH1-DsRed* Δ*arg-5* nuclei) for 6 different sectors of the same plate, represented by different colors and line styles. Likelihoods are narrowly peaked around the maximum likelihood estimator (MLEs) for each sector, and MLEs do not significantly differ between sectors of the plate. **B**: Estimated *p* (fraction of *hH1-DsRed* Δ*arg-5* nuclei) for different spore suspension proportions of the two genotypes. Changing nucleotype proportions in the starting suspension changes their abundances in the mature mycelium; pairwise differences are significant at the 𝒫 *<* 0.01 level for every experiment. Although *p* is positively correlated with the starting proportions of Δ*arg-5* spores, Δ*arg-5* nuclei are under-represented in the mature mycelium, indicating differences in fitness between the nucleotypes. **C**: Estimated proportion of *hH1-DsRed* Δ*arg-5* spores, *p*, for pre-formed heterokarya transferred to selective media. Symbols represents replicates, and shape and color represent media type. Filled symbols correspond to starting proportions of 26%:74% *hH1-DsRed* Δ*arg-5* :*hH1-eGFP* Δ*pan-2*, and unfilled symbols to 53%:47% starting proportions. *s denote significant differences at the level 𝒫 *<* 0.05. **i**: We see no significant difference in *p* between any pair of experiments that have the same spore starting proportions but different selective media. **ii**: When the same data are rearranged to compare different starting proportions, we see that starting proportions continue to control *p*. All pairwise comparisons are significant, except for media containing pantothenate (𝒫 = 0.0556). *x*-tick labels identify the starting *hH1-DsRed* Δ*arg-5* proportion, and the medium to which the heterokaryon was transferred (parenthesis). **D**: Estimated *p* when spore suspensions are germinated on selective media. Proportions are significantly different between media (** denotes 𝒫 *<* 0.05, * denotes 𝒫 *<* 0.01). In all panels, random *x*-displacements have been added to avoid overlapping points.

We first tested whether nucleotype proportions in mature mycelia are controlled by the composition of the spore suspension used to start the mycelium. We created heterokaryotic mycelia on unsupplemented minimal media using spore suspensions with different proportions of *hH1-eGFP* Δ*pan-2* and *hH1-DsRed* Δ*arg-5* nucleotypes, and analyzed the nucleotype proportions of spores produced at the edge of the plate. Nucleotype proportions in spores were linearly related to, though not equal to, the starting proportions of the two nucleotypes (Fig. 2B), over the entire range of starting proportions. In common with [26], we find no evidence that nucleotype proportions converge or even depend nonlinearly upon starting proportions. However, the ending proportions of the two nucleotypes do not match their starting proportions: for the data shown in Fig. 2B, the Δ *pan-2* nuclear population is consistently more abundant in the mature mycelium than in the spore suspension used to create the mycelium. We can advance two hypotheses for this difference: 1. mycelial level control over the abundance of the nucleotypes, since Δ *pan-2* nuclei are the heterokaryon’s only source of *arg-5* mRNAs, and during vegetative growth, *arg-5* mRNAs are 20-fold more abundant than *pan-2* mRNAs [29]; 2. the two nucleotypes have different fitnesses. Hypothesis 1 is not supported by previous experiments with these strains [26], and indeed, when we recreate (below) a subset of the starting proportions, we find Δ *pan-2* nuclei are now under-represented in the mycelium, relative to the starting spore suspension. Accordingly, we focus instead on hypothesis 2, that the two nucleotypes have different fitnesses or division rates. But are differences in division rate possible during vegetative growth, when the nuclei share a common cytoplasm? Or are they isolated to specific stages in the life-history of the heterokaryotic mycelium [17]?

To pin down where in mycelial development fitness differences between nucleotypes arise, we next studied the effect of media supplementation, which alters the degree of mutual dependence between the nucleotypes. Specifically, we added either 10*µ*L/mL calcium pantothenate, or 600*µ*L/mL L-arginine-HCl, or both, to the media on which the mycelium was growing. At this level of supplementation, wild type growth is restored in auxotrophic mutants. Thus, for example, in a *hH1-DsRed* Δ*arg-5* + *hH1-eGFP* Δ*pan-2* heterokaryon grown on calcium pantothenate supplemented medium, the nucleotypes go from being mutually to asymmetrically dependent, with the Δ*pan-2* nuclei no longer depending on Δ*arg-5* nuclei for pantothenic acid synthesis, but the Δ*arg-5* nuclei continuing to depend upon Δ*pan-2* nuclei for synthesis of arginine (Fig. 1A).

We first performed experiments where mixed spore suspensions were placed on unsupplemented minimal media to form heterokarya. After 20 hours’ growth, agar blocks containing growing hyphal tips were transferred to the supplemented medium. By transferring preformed heterokarya, we allow selection to occur only upon nucleotypes that coexist within a mycelium. We found that across all of the selective media and non-selective media, and unlike the previous experiment, *hH1-DsRed* Δ*arg-5* nuclei were over-represented in all heterokarya, relative to their starting proportions within the spore suspension, consistently for heterokarya that were started with two different initial proportions of the two nucleotypes. Thus, we conclude that there aren’t innate differences between the fitnesses of the two nucleotypes. Growth on a selective medium produced no detectable difference upon the proportion, *p* of *hH1-DsRed* Δ*arg-5* nuclei, compared to growth on unsupplemented minimal media irrespective of whether *hH1-DsRed* Δ*arg-5* nuclei were selected for, or against, by the medium (Fig. 2C i). Yet, starting proportions of the two nucleotypes continued to control the proportions seen in the grown mycelium (Fig. 2C ii).

Hypothesizing therefore that the difference between the estimated value of *p* and the starting proportion of *hH1-DsRed* Δ*arg-5* spores is due to differences in germination rates of spores or growth rate of germlings before the formation of the heterokaryon, we replicated the selective media experiment without pre-forming heterokarya by directly inoculating spore suspensions onto the selective media. Here we did see strong medium selection; including (for spore suspensions containing equal proportions of *hH1-DsRed* Δ*arg-5* and *hH1-eGFP* Δ*pan-2* nuclei) almost 4-fold greater value of *p* when spores are inoculated on arginine HCl-supplemented medium (*p* = 0.47) compared to calcium pantothenate supplemented medium (*p* = 0.12) (Fig. 2D).

## 4 Conclusions

Our work realizes, within the model ascomycete fungus *N. crassa*, the experimental framework proposed by Guido Pontecorvo when he argued that nuclear lineages within fungal heterokaryons recapitulate key features of population genetic systems. Using auxotrophic knockouts allows us to control the level of mutual dependence between nucleotypes, while fluorescent labeling of these nucleotypes enables rapid, spore-level estimates of nucleotype composition.

To emphasize the value of resolving composition at the scale of individual spores, we contrast our estimates with estimates that would be obtained if, like [26], rather than resolving individual spores, we had used e.g., flow cytometry, to gate spores by fluorescence; classifying them as homokaryotic *hH1-DsRed* Δ*arg-5* or *hH1-eGFP* Δ*pan-2* or heterokaryotic, without knowing their nuclear counts, or the correlations between nucleotypes. For example in an experiment that parallels (Fig. 2C), [26] inoculated Δ*pan* +Δ*arg* spore suspensions on medium supplemented with both pantothenate and arginine, finding that the spores collected were 63% Δ*arg-5*, 3% Δ*pan-2*, and 34% heterokaryotic. Assuming that spores are correctly classified into the three categories, we first fit these data with a binomial model, assuming *n* = 2 nuclei per spore, the least squares fit is *p* = 0.79. Assuming a greater number of nuclei per spore leads to worse fits (e.g., a 3.8-fold increase in least squares residue on the three data points if we fit for *n* = 3), as does allowing a variable number of nuclei per spore. However, accounting for the correlations between the nuclei (which reduce the likelihood that a spore with *n* = 2 nuclei is heterokaryotic), via the beta-binomial distribution, and imposing a pairwise correlation between nuclei,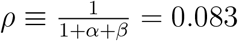, comparable to our own experiments, then the minimum residue occurs for *n* = 8, and has *p* = 0.65, while constraining *n* = 2, produces even smaller bias toward Δ*arg* : *p* = 0.59.

In contrast to fluorotyping, qPCR methods can provide unambiguous readouts of the nucleotype proportions within groups of spores via ratios of gene markers, and can also assay the nucleotype composition of other fungal cell types (such as the hyphae themselves, or the specialized tissues associated with sexual reproduction) [17]. Although sequencing-based methods can’t assay diversity at the level of individual hyphae, *N. crassa*’s hyphal architecture is organized to mix genetically diverse nuclei, so that hyphal heterogeneity may be small [6, 25]. However, fluorotyping of individual spores directly shows the interplay between number of nuclei per spore and nucleotype diversity. For example, although we focused upon the parameter *p* by measuring the proportion of *hH1-DsRed* Δ*arg-5* nuclei across all sampled spores, the proportion of spores that contain any *hH1-DsRed* Δ*arg-5* nucleus is, itself, an important fitness readout, since it may be intepreted as a proxy for the proportion of asexually generated offspring mycelia in which the *hH1-DsRed* Δ*arg-5* nucleotype would be present. Even when we found high skews toward one nucleotype, a large number of spores, particularly spores with large numbers of nuclei, contained at least one nucleus with each nucleotype. Nuclear number per spore depends on external conditions [30], and potentially allows the proportion of heterokaryotic offspring to be controlled. Indeed, although many studies have illuminated how, in the distantly related Glomeromycota, asexual megaspores enable heterokaryotic mycelia to transmit genetically diverse nuclei [31], the parallel question of the extent to which *N. crassa* or other filamentous Ascomycota might manipulate asexual spore size to maximize their genetic diversity remains unanswered. Correlations between nuclei in spores that were parameterized by our beta binomial model, and meant that spores, even ones with many nuclei, are less likely to be genetically diverse than one would expect based on the number of nuclei they contain. The likely mechanism for diversity loss is imperfect mixing of apical nuclei during conidiogenesis. Bulk flow of cytoplasm can mix nuclei and organelles in the interior of the mycelium, but is less effective for mixing at hyphal tips, including potentially in the aerial hyphae that produce *N. crassa*’s conidiospores [6, 32]. But nuclei in *Neurospora* also have microtubule-associated motility [33], which is known in another ascomycete, *Ashbya gossypii*, to mix nuclear lineages [34, 35]. Thus, by controlling nuclear motions within aerial hyphae, the heterokaryotic fungus may be able to control the nucleotypic diversity of its spores.

A new image analysis pipeline for fluorotyping spores forms the backbone of this paper’s scientific contributions. With this new approach, we resurface a question posed by Pontecorvo [11]: when does selection of nucleotypes occur during the growth of heterokaryotic *N. crassa* mycelia? Comparing Fig. 2C and 2D, we find a reaffirmation of Pontecorvo’s prediction: that media can select for one of the components of the heterokaryon if exposure occurs before the establishment of a single linked mycelium (Fig. 2D), but when a preformed heterokaryotic mycelium is transferred to a selective medium, then the sharing of resources between nuclei seems to preclude any changes in nucleotype ratios (Fig. 2C). We conclude that both arginine and pantothenic acid are freely shared between nuclei across the cytoplasm [32].

Continuing to support both nucleotypes, even on media in which one nucleotype could have grown independently may serve as a form of *bet-hedging*, in case the fungus encounters an environment for which the genes present in the other nucleotype are adapted [26, 12]. However, our results emphasize that, although *N. crassa*’s conidia have a known role in vegetative reproduction and dispersal [36], they also can function as genetic bottlenecks: winnowing down the number of nucleotypes present in a single *N. crassa* mycelium [3] to a smaller sample, and subjecting these samples to competition against each other, before an offspring mycelium is created. Tolerance of less fit nucleotypes exposes the heterokaryotic mycelium to a form of Muller’s ratchet, or accumulation of deleterious mutations. Although the importance of unicellular bottlenecks has been emphasized across multicellular organisms [37, 5], including in *N. crassa* [38], our work invites study of the plasticity of *Neurospora*’s pre-heterokaryon formation phase: even when there is strong selection for a nucleotype that is adapted to the spore germination medium, the less-fit nucleotype was still present in all mycelia, and the degree to which nuclei from these spores are admitted into the heterokaryon depends on the (potentially controllable) relative timings of conidial germination, nuclear division, and formation of the conidial anastomosis tubes through which spores fuse [39]. In this sense, to Pontecorvo’s analogy between the nucleotypes interacting in a heterokaryon, and the population genetics of interacting organisms, we can add a second analogy between the timings of germination and fusion events during heterokaryon formation and the timing of germline segregation during embryo development [5]. Within embryos, segregation of germline cells limits the impact of selfish somatic cell mutations, and resolves conflicts between cell lineages within individual organisms [5]. But in animals, including mammals, in which germ lines are established epigenetically, the timing of segregation may also be plastic, and its timing directly regulates how much competition can occur between cell lineages [5]. Selective forces that act on heterokaryotic groups of spores but not upon heterokaryotic mycelia create an analogous context in which mechanisms such as transcriptional suppression of the nuclei present in the spore and/or carrying over of mRNAs from the originating mycelium could play an analogous role to the maternal mRNAs present in multicellular embryos [40].

The influence of the timings of spore germination and fusion also likely determines why different experiments, set up from different spore suspensions, have limited reproducibility. For example, the data used to generate Fig. 2B and Fig. 2C both include data points in which the starting spore compositions were approximation 50% of each of the two strains, and spores were plated onto (heterokaryon-inducing) minimal media. Yet, the estimated *p*, while similar between replicates derived from the same slant tubes at the same time, are significantly different between ‘equivalent’ experiments that are performed using spore suspensions derived from different slant tubes at different times (*p* = 0.63 for Fig. 2C and *p* = 0.24 for 2B): in one experiment the heterokaryon proportions skew toward *hH1-DsRed* Δ*arg-5* and in the other, they skew toward *hH1-eGFP* Δ*pan-2* . We attribute the different outcomes to the sensitivity of the heterokarya to the conditions of pre-formation. The condition of the fungus plays a role in the germination rates of its spores [41]. Although we sought to minimize differences in spore condition by using spores of similar (and matched) ages between experiments, the effect size remains as large as inoculating spores onto selective media. What other factors led to these differences? As our image analysis platform is able to establish a profile for each individual spore, other potential determinants of spore-condition, including the spore size, nucleocytoplasmic ratio and lipid content can be investigated in future study.

## Acknowledgement

We acknowledge financial support from NSF (Grant RoL-1840273).We thank Samuel Christensen for helpful discussions.

## References

[1] Alexander P Mela, Adriana M Rico-Ramírez, and N Louise Glass. Syncytia in fungi. Cells, 9(10):2255, 2020.

[2] Marcus Roper, Chris Ellison, John W Taylor, and N Louise Glass. Nuclear and genome dynamics in multinucleate ascomycete fungi. Current Biology, 21(18):R786–R793, 2011.

[3] Ramesh Maheshwari. Nuclear behavior in fungal hyphae. FEMS Microbiology Letters, 249(1):7–14, 2005.

[4] Amy S Gladfelter, A Katrin Hungerbuehler, and Peter Philippsen. Asynchronous nuclear division cycles in multinucleated cells. The Journal of Cell Biology, 172(3):347–362, 2006.

[5] Leo W Buss. The evolution of individuality. Princeton University Press, 2014.

[6] Marcus Roper, Anna Simonin, Patrick C Hickey, Abby Leeder, and N Louise Glass. Nuclear dynamics in a fungal chimera. Proceedings of the National Academy of Sciences, 110(32):12875–12880, 2013.

[7] James B Anderson, Johann N Bruhn, Dahlia Kasimer, Hao Wang, Nicolas Rodrigue, and Myron L Smith. Clonal evolution and genome stability in a 2500-year-old fungal individual. Proceedings of the Royal Society B, 285(1893):20182233, 2018.

[8] James K Hane, Jonathan P Anderson, Angela H Williams, Jana Sperschneider, and Karam B Singh. Genome sequencing and comparative genomics of the broad host-range pathogen rhizoctonia solani ag8. PLoS Genetics, 10(5):e1004281, 2014.

[9] Asen Daskalov, Jens Heller, Stephanie Herzog, André Fleißner, and N Louise Glass. Molecular mechanisms regulating cell fusion and heterokaryon formation in filamentous fungi. Microbiology Spectrum, 5(2):10–1128, 2017.

[10] Timothy L Friesen, Eva H Stukenbrock, Zhaohui Liu, Steven Meinhardt, Hua Ling, Justin D Faris, Jack B Rasmussen, Peter S Solomon, Bruce A McDonald, and Richard P Oliver. Emergence of a new disease as a result of interspecific virulence gene transfer. Nature Genetics, 38(8):953–956, 2006.

[11] G Pontecorvo. Genetic systems based on hetero caryosis. In Cold Spring Harbor Symposia on Quantitative Biology, volume 11, pages 193–201. Cold Spring Harbor Laboratory Press, 1946.

[12] CE Caten. The mutable and treacherous tribe revisited. Plant Pathology, 45(1), 1996.

[13] László G Nagy, Torda Varga, Árpád Csernetics, and Máté Virágh. Fungi took a unique evolutionary route to multicellularity: Seven key challenges for fungal multicellular life. Fungal Biology Reviews, 34(4):151–169, 2020.

[14] Alan DM Rayner. The challenge of the individualistic mycelium. Mycologia, 83(1):48– 71, 1991.

[15] Timothy Y James, Jan Stenlid, Åke Olson, and Hanna Johannesson. Evolutionary significance of imbalanced nuclear ratios within heterokaryons of the basidiomycete fungus Heterobasidion parviporum. Evolution, 62(9):2279–2296, 2008.

[16] H Rees and JL Jinks. The mechanism of variation in Penicillium heterokaryons. Proceedings of the Royal Society of London. Series B-Biological Sciences, 140(898):100–106, 1952.

[17] Cécile Meunier, Sara Hosseini, Nahid Heidari, Zaywa Maryush, and Hanna Johannesson. Multilevel selection in the filamentous ascomycete Neurospora tetrasperma. The American Naturalist, 191(3):290–305, 2018.

[18] Cécile Meunier, Iulia Darolti, Johan Reimegård, Judith E Mank, and Hanna Johannesson. Nuclear-specific gene expression in heterokaryons of the filamentous ascomycete neurospora tetrasperma. Proceedings of the Royal Society B, 289(1980):20220971, 2022.

[19] Eric Bastiaans, Alfons JM Debets, and Duur K Aanen. Experimental evolution reveals that high relatedness protects multicellular cooperation from cheaters. Nature Communications, 7(1):11435, 2016.

[20] Anne Pringle and John W Taylor. The fitness of filamentous fungi. Trends in Microbiology, 10(10):474–481, 2002.

[21] Seth Rakoff-Nahoum, Kevin R Foster, and Laurie E Comstock. The evolution of cooperation within the gut microbiota. Nature, 533(7602):255–259, 2016.

[22] Wentao Kong, David R Meldgin, James J Collins, and Ting Lu. Designing microbial consortia with defined social interactions. Nature Chemical Biology, 14(8):821–829, 2018.

[23] Jeremy J Minty, Marc E Singer, Scott A Scholz, Chang-Hoon Bae, Jung-Ho Ahn, Clifton E Foster, James C Liao, and Xiaoxia Nina Lin. Design and characterization of synthetic fungal-bacterial consortia for direct production of isobutanol from cellulosic biomass. Proceedings of the National Academy of Sciences, 110(36):14592–14597, 2013.

[24] Jeff Gore, Hyun Youk, and Alexander Van Oudenaarden. Snowdrift game dynamics and facultative cheating in yeast. Nature, 459(7244):253–256, 2009.

[25] Linda Ma, Boya Song, Thomas Curran, Nhu Phong, Emilie Dressaire, and Marcus Roper. Defining individual size in the model filamentous fungus Neurospora crassa. Proceedings of the Royal Society B: Biological Sciences, 283(1826):20152470, 2016.

[26] Alexander P Mela and N Louise Glass. Permissiveness and competition within and between Neurospora crassa syncytia. Genetics, 224(4):iyad112, 2023.

[27] Michael Freitag and Eric U Selker. Expression and visualization of red fluorescent protein (RFP) in Neurospora crassa. Fungal Genetics Reports, 52(1):14–17, 2005.

[28] Henry J Vogel. A convenient growth medium for neurospora (medium n). Microbial Genet. Bull., 13:42–43, 1956.

[29] Huiquan Liu, Yang Li, Daipeng Chen, Zhaomei Qi, Qinhu Wang, Jianhua Wang, Cong Jiang, and Jin-Rong Xu. A-to-I RNA editing is developmentally regulated and generally adaptive for sexual reproduction in Neurospora crassa. Proceedings of the National Academy of Sciences, 114(37):E7756–E7765, 2017.

[30] Charles Huebschman. A method for varying the average number of nuclei in the conidia of Neurospora crassa. Mycologia, 44(5):599–604, 1952.

[31] Jean-Luc Jany and Teresa E Pawlowska. Multinucleate spores contribute to evolutionary longevity of asexual glomeromycota. The American Naturalist, 175(4):424–435, 2010.

[32] Marcus Roper, ChangHwan Lee, Patrick C Hickey, and Amy S Gladfelter. Life as a moving fluid: fate of cytoplasmic macromolecules in dynamic fungal syncytia. Current Opinion in Microbiology, 26:116–122, 2015.

[33] Silvia L Ramos-García, Robert W Roberson, Michael Freitag, Salomón Bartnicki-García, and Rosa R Mouriño-Pérez. Cytoplasmic bulk flow propels nuclei in mature hyphae of Neurospora crassa. Eukaryotic Cell, 8(12):1880–1890, 2009.

[34] Sandrine Grava and Peter Philippsen. Dynamics of multiple nuclei in Ashbya gossypii hyphae depend on the control of cytoplasmic microtubules length by Bik1, Kip2, Kip3, and not on a capture/shrinkage mechanism. Molecular Biology of the Cell, 21(21):3680– 3692, 2010.

[35] Cori A Anderson, Umut Eser, Therese Korndorf, Mark E Borsuk, Jan M Skotheim, and Amy S Gladfelter. Nuclear repulsion enables division autonomy in a single cytoplasm. Current Biology, 23(20):1999–2010, 2013.

[36] Carmen Ruger-Herreros and Luis M Corrochano. Conidiation in Neurospora crassa: vegetative reproduction by a model fungus. International Microbiology, 23(1):97–105, 2020.

[37] Richard K Grosberg and Richard R Strathmann. One cell, two cell, red cell, blue cell: the persistence of a unicellular stage in multicellular life histories. Trends in Ecology & Evolution, 13(3):112–116, 1998.

[38] Eric Bastiaans, Duur K Aanen, Afons JM Debets, Rolf F Hoekstra, Bram Lestrade, and Marc FPM Maas. Regular bottlenecks and restrictions to somatic fusion prevent the accumulation of mitochondrial defects in neurospora. Philosophical Transactions of the Royal Society B: Biological Sciences, 369(1646):20130448, 2014.

[39] M Gabriela Roca, Nick D Read, and Alan E Wheals. Conidial anastomosis tubes in filamentous fungi. FEMS Microbiology Letters, 249(2):191–198, 2005.

[40] Leah Vardy and Terry L Orr-Weaver. Regulating translation of maternal messages: multiple repression mechanisms. Trends in Cell Biology, 17(11):547–554, 2007.

[41] S Earl Kang, Brandi N Celia, Douda Bensasson, and Michelle Momany. Sporulation environment drives phenotypic variation in the pathogen Aspergillus fumigatus. G3, 11(8):jkab208, 2021.

