## Supplemental Information for "Life history stage-dependent nuclear selection in chimeric fungi"

### 1 Description

In the Supplementary Information we describe the image anaylsis pipeline that was created to fluorotype spores (classify them by their fluorescence), and to estimate the nucleotype proportions in the heterokaryotic mycelium from which spores were harvested. Our analytical method consists of two image analysis steps, followed by two data analysis steps.

### 2 Steps 1 & 2: Detection of Nuclei and Measurement of Fluorescence Intensities

The gray-scale transmitted light images were binarized with an adaptive threshold based on the local mean intensity of each pixel, which gave the rough outlines of the spores. Frequently, some spores clustered together, so that segmentation was required. After filtering out the local minimum with low signal to noise ratio in the gray-scale image, a distance transform [1] was applied, which determined the distance from the foreground pixel to the nearest background pixel. The watershed transformation [2] was then used to partition clusters of spores. Finally, objects that were close to the border of the image, smaller than 18.75 $\mu m^2$, or larger than 125 $\mu m^2$ were removed.

We used template matching [3] to identify potential nuclei in the corresponding fluorescence images for heterokaryotic spores [4]. We manually chose a template showing a typical nucleus and resized it to account for nuclei of different sizes. We then applied template matching based on the relative intensity to the fluorescence images in DsRed and EGFP channel separately. We consider all local maxima of the template match image, and call these *candidate nuclei*. For the candidates in DsRed channel, we filtered out all spots whose fluorescence intensity were less than either: $\frac{1}{2}$ of the mean fluorescence intensity of the detected spots, or $\frac{1}{10}$ of the maximum intensity. For the spots in EGFP channel, we filtered out spots with intensities less than the greater of $\frac{1}{5}$ of the maximum intensity of the detected spots and 400. The criteria for *candidate nuclei* are different in DsRed and GFP channel due to the variation in level of expression. We checked by eye that this method agreed with by eye segmentation of nuclei in all experimental conditions Finally, we overlaid the nuclei positions in EGFP and DsRed channels. For each spore, the number of nuclei, $N_i$ was found from whichever is greater, the number of spots in the DsRed channel, or the number in the GFP channel. If a spore contained nuclei with only one genotype it is tagged as homokaryotic DsRed/EGFP, otherwise, the total (summed) intensity of all spots across the GFP and DsRed channels were recorded as variables $G_i$ and

$R_i$, respectively.

#### 3 Steps 3 & 4: Inferring nucleotype proportions

With the data available, we estimate $p$, the probability that a detected nucleus is *hH1-DsRed* $\Delta$*arg-5* . Previous studies have shown that the different nuclei packaged within the same conidium have correlated nucleotypes [5, 6]: that is, the mycelium produces more homokaryotic conidia than would be expected if nuclei were independently drawn into the spore from over the entire mycelium, likely as result to imperfect mixing of nucleotypes at the scale of single hyphae [7]. Our method allows us to collect genotype data for mono- and bi- nucleate spores, and to directly affirm that the nucleotypes are correlated in a single spore. We model these correlations and the subsequent overdispersion of nucleotypes within multinucleate spores via a beta binomial distribution [8]. That is, we assume that in a spore containing $n$ nuclei, the number that are *hH1-DsRed* $\Delta$*arg-5* is Binomial$(n, P)$ distributed, where for each spore, the $P$ are independent Beta$(\alpha, \beta)$ random variables, and the parameters $\alpha$ and $\beta$, assumed constant across all spores from a single plate, are estimated via likelihood maximization. Throughout, we report the mean proportion of *hH1-DsRed* $\Delta$*arg-5* nuclei across all spores, i.e., $p \equiv \langle P \rangle = \frac{\alpha}{\alpha+\beta}$.

To estimate $\alpha$ and $\beta$ we divide spores into two classes: homokaryotic (which can be definitively genotyped, since they have significant fluorescence only in one channel) and heterokaryotic, for which the number of nuclei of each type becomes a hidden fitting variable, and each with each possible partitioning of nuclei into nucleotypes is assigned a likelihood based on the observed total fluorescent intensities $(R, G)$ of the spore. (Because of protein sharing between nuclei, it is not possible to genotype individual nuclei based on their fluorescence, but we assume that the total DsRed and GFP intensities of the spore reflect the number of each nucleotype present).

For each plate, we estimate the contribution of a single *hH1-DsRed* $\Delta$*arg-5* or *hH1-eGFP* $\Delta$*pan-2* nucleus to the fluorescence intensity of the spore that it contain, by first identifying all mononucleate spores, each of which must be either an *hH1-eGFP* $\Delta$*pan-2* or *hH1-DsRed* $\Delta$*arg-5* homokaryon. We used the data of those spores to approximate the DsRed intensity of a single *hH1-DsRed* $\Delta$*arg-5* nucleus, and GFP intensity of a single *hH1-eGFP* $\Delta$*pan-2* nucleus, respectively $I_{rr} \sim N(\mu_{rr}, \sigma_{rr})$ and $I_{gg} \sim N(\mu_{gg}, \sigma_{gg})$. Mononucleate spores also had fluorescence in the other channel (that is, a spore containing only one *hH1-DsRed* $\Delta$*arg-5* nucleus, will also have some fluorescence in the GFP channel), likely due to incomplete turnover of proteins that were packaged in the spore at its formation. We call the cross-channel fluorescences

$\mu_{gr}$ for the DsRed fluorescence of a *hH1-eGFP* $\Delta$*pan-2* spore and $\mu_{rg}$ for the GFP fluorescence of a *hH1-DsRed* $\Delta$*arg-5* spore, and treat both as constants (since the variation in intensity is so much less than the in-channel fluorescences $I_{rr}$ and $I_{gg}$. In most cases we found enough single spores of both nucleotypes that we could estimate all 6 parameters, and validate the hypothesis of normal distributions (Fig. 1). Where a single plate did not contain enough mononucleate spores of both types to estimate all parameters, we aggregated all of the mononucleate spores from the same experimental condition.

Under these assumptions, the DsRed and GFP intensities of a spore containing $N$ nuclei, of which $k$ are *hH1-DsRed* $\Delta$*arg-5* is:

$$(R|k) \sim N(k\mu_{rr} + (N-k)\mu_{gr}, k\sigma_{rr}^2) \tag{1}$$

$$(G|k) \sim N(k\mu_{rg} + (N-k)\mu_{gg}, (N-k)\sigma_{gg}^2) \tag{2}$$

For each spore $i$, we calculate the likelihood for the observed variables ($N_i$, $G_i$, $R_i$):

$$\begin{aligned} \mathcal{L}(R_i, G_i|\alpha,\beta) = & \sum_{k=1}^{N_i-1} \mathcal{L}(R_i, G_i|k)\mathcal{L}(k|\alpha,\beta) \\ = & \sum_{k=1}^{N_i-1} \exp\left[-\frac{(R_i-(k\mu_{rr}+(N_i-k)\mu_{gr}))^2}{2k\sigma_{rr}^2} - \frac{(G_i-(k\mu_{rg}+(N_i-k)\mu_{gg}))^2}{2(N_i-k)\sigma_{gg}^2}\right] \times \\ & \frac{1}{2\pi\sqrt{k(N_i-k)\sigma_{rr}^2\sigma_{gg}^2}}\binom{N_i}{k}\frac{B(\alpha+k,N_i-k+\beta)}{B(\alpha,\beta)} \ . \end{aligned} \tag{3}$$

For each experimental condition, we estimate the likelihood maximizing parameters $\alpha$ and $\beta$ by defining 100 evenly spaced values between 0.01 and 10 for both parameters, and by computing the joint log-likelihood across all spores for each combination of $(\alpha, \beta)$ (Fig. 2). In each evaluated case there us a local maximum value of the log-likelihood, for which we report the parameters $(\alpha, \beta)$. Importantly, although the likelihood landscape is quite flat along a line $\alpha/\beta$ = constant (Fig. 2), obtaining the value of $\alpha/\beta$, allows for computation of $p = \alpha/(\alpha+\beta)$; in other words, the mean proportion of *hH1-DsRed* $\Delta$*arg-5* nuclei can be readily calculated, though parameters associated with the correlated packaging of nuclei into spores are harder to estimate from the given data. We report posterior uncertainties in $p$ by plotting the likelihood (normalized so the maximum likelihood is 1), for a quarter-circular arc of parameters $(\alpha, \beta)$ (which meets the line $p$ = constant at right angles).

Throughout this work, non-parametric permutation tests [9] were used to quantify the difference in $p$-estimates between different conditions.

For each replicate (plate), we combine data from all 20+ images ($\simeq$ 1500 - 6000 spores) to make a single estimate of $p$. To estimate confidence bounds upon $p$, we subdivided the sets of images into either 5 or 6 groups (i.e., 4-5 images per group), and

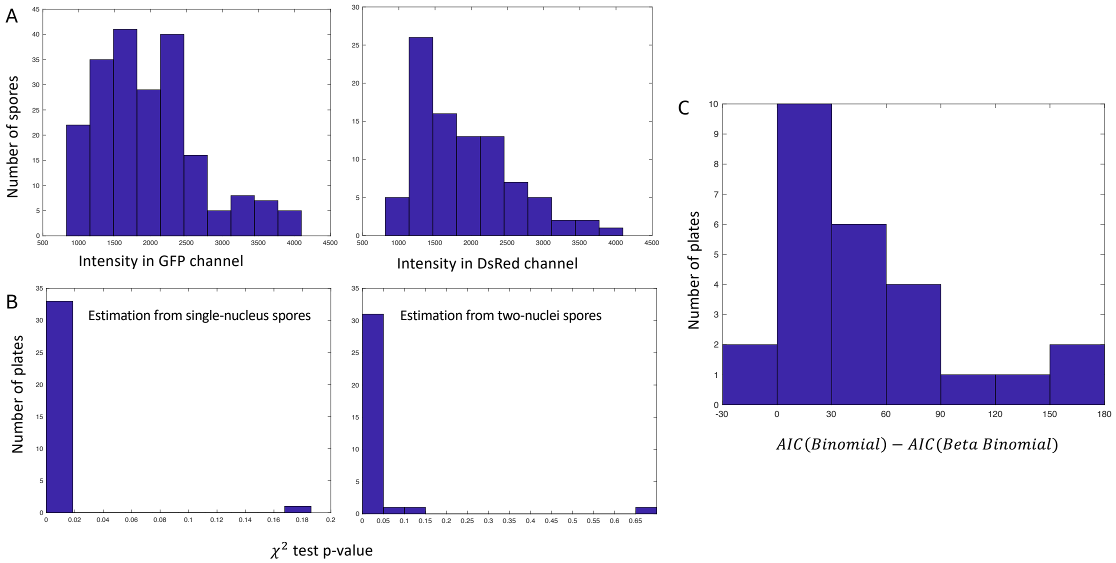


Figure 1: **Spore fluorescence is converted into nucleotype intensities via empirically parameterized models.** **A**: Typical same channel fluorescence intensity distributions for $I_{gg}$ (GFP intensity of a *hH1-eGFP* $\Delta$*pan-2* nucleus; left) and $I_{rr}$ (DsRed intensity of a *hH1-DsRed* $\Delta$*arg-5* nucleus; right), estimated from mononucleate conidia. We model both distributions as approximately normal. A one-sample Kolmogrov-Smirnov normality test compares $\mathcal{P}$-values 0.1846 for $I_{gg}$ and 0.0972 for $I_{rr}$. **B**: Distribution of $\mathcal{P}$-values for $\chi^2$ goodness of fit tests for 34 experiments, assuming nuclei are independently packaged in mono- and dikaryotic spores. Most of the $\mathcal{P}$-value are less than 0.05 against independence. **C**: Difference in AIC scores for Beta-binomial and binomial fits to the same data shown fitted for panel B. Positive values indicate better fit for beta-binomial distribution than binomial, even when additional fitting parameter is accounted for.

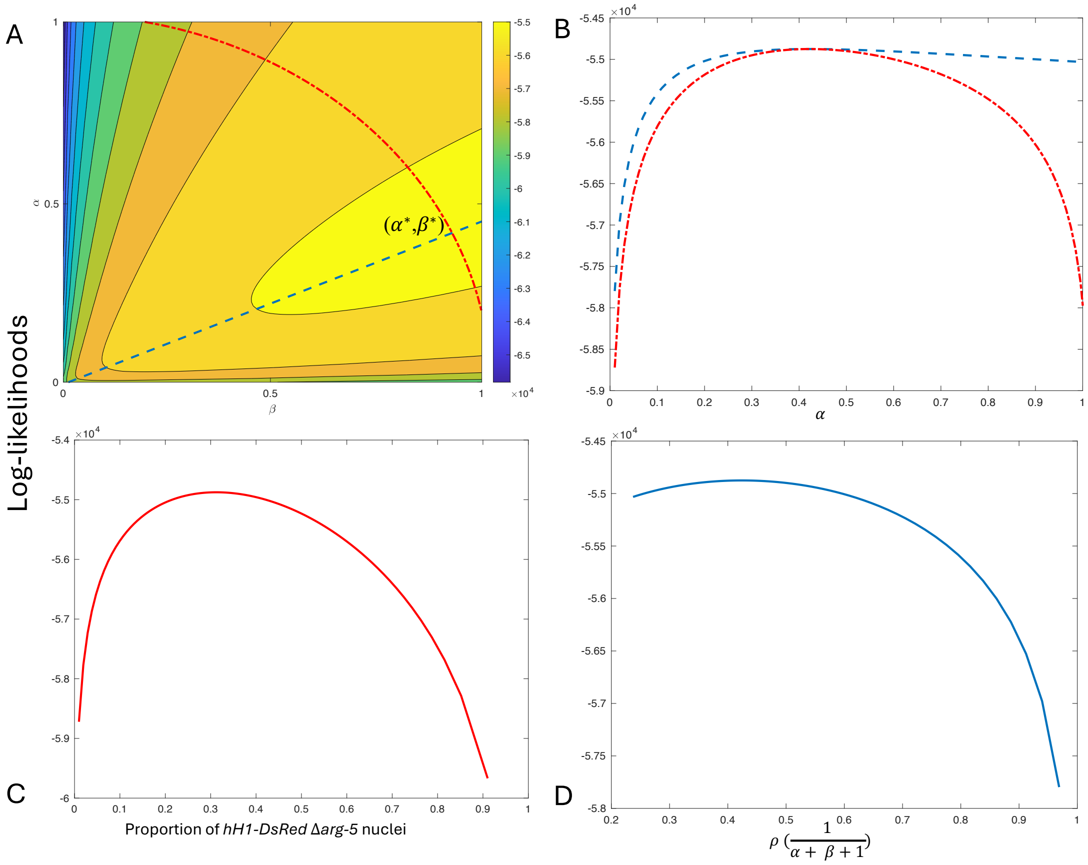


Figure 2: **Beta-binomial fits give maximum likelihood values for the parameters $\alpha$ and $\beta$, which can be interpreted in terms of proportion of *hH1-DsRed* $\Delta$*arg-5* nuclei, and nuclear-pair-correlations within single spores.** **A**: joint log-likelihoods across all spores in a single condition (here a plate inoculated with a 59:41 ratio of *hH1-DsRed* $\Delta$*arg-5* to *hH1-eGFP* $\Delta$*pan-2* spores)for different $\alpha, \beta$ values, which is maximized at $(\alpha^*, \beta^*)$. Blue dashed line: where $\frac{\alpha}{\alpha+\beta} = \frac{\alpha^*}{\alpha^*+\beta^*}$.Red dashed line: where $\alpha^2 + \beta^2 = \alpha^{*2} + \beta^{*2}$. **B**: the log-likelihoods along the red dashed line and the blue dashed line for different $\alpha$. **C**: the log-likelihoods along the red dashed line with respect to the proportion of *hH1-DsRed* $\Delta$*arg-5* nuclei. **D**: the log-likelihoods along the blue dashed line plotted against the pair-wise correlation of the beta-binomial distribution.

reran our $p$-estimate analysis on the images in each group separately. We calculate the standard error of these 4-5 different $p$-estimates. Since the subdivided images might not contain enough homokaryotic spores for reliable estimation of $p_{rr}$ and $p_{gg}$, we used the $p_{rr}$ and $p_{gg}$ built from the complete data for those subgroup estimations.

### References


[1] Calvin R Maurer, Rensheng Qi, and Vijay Raghavan. A linear time algorithm for computing exact euclidean distance transforms of binary images in arbitrary dimensions. *IEEE Transactions on Pattern Analysis and Machine Intelligence*, 25(2):265–270, 2003.

[2] Fernand Meyer. Topographic distance and watershed lines. *Signal processing*, 38(1):113–125, 1994.

[3] Dirk-Jan Kroon. Fast/Robust Template Matching. https://www.mathworks.com/matlabcentral/fileexchange/24925-fast-robust-template-matching, 2011. Online; accessed February 2, 2022.

[4] Linda Ma, Boya Song, Thomas Curran, Nhu Phong, Emilie Dressaire, and Marcus Roper. Defining individual size in the model filamentous fungus Neurospora crassa. *Proceedings of the Royal Society B: Biological Sciences*, 283(1826):20152470, 2016.

[5] Timothy Prout, Charles Huebschman, Howard Levene, and Francis J Ryan. The proportions of nuclear types in Neurospora heterocaryons as determined by plating conidia. *Genetics*, 38(5):518, 1953.

[6] KC Atwood and Frank Mukai. Nuclear distribution in conidia of Neurospora heterokaryons. *Genetics*, 40(4):438, 1955.

[7] Marcus Roper, Anna Simonin, Patrick C Hickey, Abby Leeder, and N Louise Glass. Nuclear dynamics in a fungal chimera. *Proceedings of the National Academy of Sciences*, 110(32):12875–12880, 2013.

[8] John Gordon Skellam. A probability distribution derived from the binomial distribution by regarding the probability of success as variable between the sets of trials. *Journal of the Royal Statistical Society. Series B (Methodological)*, 10(2):257–261, 1948.

[9] Laurens R Krol. Permutation test. https://www.mathworks.com/matlabcentral/fileexchange/63276-permutation-test, 2024. [Online; accessed June 5, 2025].
